# Epilepsy and premature mortality driven by inhibitory neuron dysfunction in a mouse model of *SCN1A* gain-of-function neurodevelopmental disorder

**DOI:** 10.64898/2026.08.04.742893

**Authors:** Sophie F. Hill, Zachary P. Rosenthal, Ethan M. Goldberg

## Abstract

The gene most commonly implicated in epilepsy, *SCN1A*, encodes the neuronal voltage-gated sodium channel subunit NaV1.1. *SCN1A* variants that reduce sodium current (“loss of function” variants) cause Dravet syndrome, a neurodevelopmental disorder defined by treatment-resistant temperature-sensitive epilepsy with onset at/around 5 months of age, developmental delay/intellectual disability, and features of or formal diagnosis autism. However, an emerging group of variants cause “gain of function” (GoF) effects on NaV1.1 and result in a distinct presentation with earlier onset than Dravet syndrome and prominent movement disorder but without temperature sensitivity. We developed the first mouse model of *SCN1A* GoF epilepsy with heterozygous Cre-dependent expression of the recurrent patient variant *Scn1a*-p.R1636Q. Global expression of this variant causes premature mortality in 100% (64/64) of mutant mice between postnatal day 12-18 due to spontaneous, convulsive seizures. Activation of the mutant allele in parvalbumin interneurons (*Dlx5/6-Cre* or *PV-Cre*), but not excitatory neurons (*Slc17a7-Cre*) or other interneuron subtypes (*VIP-Cre* or *Sst-Cre*), recapitulates the premature mortality and epilepsy phenotypes. Treatment of *Scn1a*-p.R1636Q mutant mice with the sodium channel blocker GS967 markedly prolongs lifespan. This work is the first study of *SCN1A* GoF epilepsy in a preclinical model *in vivo*. Further investigation in the *Scn1a*^*flox(R1636Q)*^mouse will yield new mechanistic insights into disease mechanisms to drive advances in the treatment of *SCN1A* GoF epilepsy.

## Introduction

Variants in *SCN1A*, which encodes the voltage-gated sodium channel subunit Na_V_1.1, are the most common genetic cause of epilepsy (1–3). In 2000, the first pathogenic *SCN1A* variants were discovered in two families with generalized (now called genetic) epilepsy with febrile seizures plus (GEFS+). GEFS+ is on the milder end of the *SCN1A* phenotypic spectrum and is characterized by febrile seizures that continue beyond age 5-6 years and may be accompanied by typically mild epilepsy that often resolves in adulthood (4, 5). The following year, missense, nonsense, and frameshift variants in *SCN1A* were discovered in 7 patients with Dravet syndrome, now known to be the most common genetic epilepsy (6, 7). Dravet syndrome is a neurodevelopmental disorder characterized by symptom onset at/around 5 months of age, treatment-resistant epilepsy with prominent temperature sensitivity, developmental delay/intellectual disability, and features of autism spectrum disorder with a formal diagnosis of autism in up to 70% of cases (8, 9).

The presence of *SCN1A* nonsense and frameshift variants in Dravet syndrome patients suggested that the disease mechanism was haploinsufficiency, a surprising result in 2001 given the role of sodium channels in promoting action potential firing. Indeed, in the initial report of GEFS+ patients with *SCN1A* missense variants, the authors speculated that the variants might delay channel inactivation (5). Subsequent functional characterization, however, disproved this original hypothesis and confirmed that Dravet syndrome and GEFS+ associated *SCN1A* variants reduce the sodium current passed by Na_V_1.1 and can be classified as “loss-of-function” (LoF) (10, 11).

The development of *Scn1a*^*+/-*^ mouse models clarified the pathogenic mechanism driving Dravet syndrome. In fact, Na_V_1.1 is the primary sodium channel α-subunit expressed in GABAergic inhibitory neurons, with particularly high expression in fast-spiking parvalbumin-expressing interneurons (PV+INs) (12–14). Currently, the leading mechanistic hypothesis of Dravet syndrome posits that loss of *SCN1A* impairs action potential generation and limits the firing frequency of PV+INs during development, leading to disinhibition and overall network hyperexcitability (13, 15).

However, a growing group of *SCN1A* variants cause elevated Na_v_1.1 current (“gain-of-function”, GoF), usually by impairing channel inactivation and/or increasing persistent sodium current (6, 16–18). *SCN1A* GoF variants cause two strikingly different phenotypes. Some variants result in familial hemiplegic migraine type 3 (FHM3), an inherited migraine condition with hemiplegic aura, sometimes accompanied by epilepsy but typically in the context of normal development and intellectual capacity (17).

Other *SCN1A* GoF variants cause an even more severe neurodevelopmental disorder than Dravet syndrome (often termed early infantile epileptic encephalopathy with movement disorder, EIDEE-MD), with onset often in the first 3 months of life (and in the first days or weeks of life in many cases), multiple seizure types, prominent movement disorder, and severe to profound intellectual disability (10, 11). Some individuals with especially early seizure onset also exhibit arthrogryposis multiplex congenita, suggesting a particularly severe movement disorder with onset *in utero* (termed neonatal developmental and epileptic encephalopathy with movement disorder and arthrogryposis, NDEEMA) (11, 19). It is not yet clear why some variants cause a neurodevelopmental disorder with onset in infancy and others cause a relatively less severe migraine disorder in the setting of normal cognitive function (10, 11, 17). *SCN1A* GoF epilepsy in particular is poorly understood, in part due to a lack of experimental model systems.

Here, we report the development and characterization of the first mouse model of *SCN1A* GoF epilepsy, which harbors the recurrent p.R1636Q variant (10). Global expression of the conditional variant via cross to *Actb-Cre* causes epilepsy and premature mortality by postnatal day (P)18. This phenotype was largely recapitulated by expressing the variant in inhibitory neuron progenitors (*Dlx5/6-Cre*) and partially by postnatal expression in parvalbumin interneurons (*PV-Cre*), but not by expressing the variant in excitatory neurons (*Slc17a7-Cre*), somatostatin interneurons (*SstCre*), or vasoactive intestinal peptide interneurons (*Vip-Cre*). Treatment with the sodium channel blocker GS967 protected mice against premature death. This work further supports dysfunction of GABAergic interneurons as the locus of *SCN1A* spectrum disorders, as well as an early developmental role of GABAergic inhibitory interneuron dysfunction in the pathogenesis of *SCN1A* GoF epilepsy. The *Scn1a*^*flox(R1636Q)*^ mouse recapitulates many aspects of the human *SCN1A* GoF epilepsy phenotype, suggesting this model will be useful in advancing the understanding of pathogenic mechanisms and development of novel therapies.

## Methods

### Mice

All experiments were approved by the Institutional Animal Care and Use Committee at the Children’s Hospital of Philadelphia (CHOP) and conducted in accordance with the ethical guidelines of the National Institutes of Health.

Male and female mice were used for all experiments in roughly equal proportions. All mice were on a 100% C57/Bl6J genetic background. Litters were weaned between P21-28, and males and females were subsequently housed separately with up to 5 animals per cage. Animals were maintained on a 12/12h light/dark cycle with ad libitum access to food and water.

Establishment of the *Scn1a*^*flox(R1636Q)*^ mouse is described below. Experimental mice were generated by crossing *Scn1a*^*flox(R1636Q)*^ heterozygous or homozygous male or female mice to a Cre driver line. Cre lines used were: *Actb-Cre* (global expression, RRID:IMSR_JAX:019099), *Slc17a7-Cre* (excitatory neurons, RRID:IMSR_JAX:037512), *Dlx5/6-Cre* (inhibitory neuron progenitors, RRID:IMSR_JAX:008199), *PV-Cre* (mature parvalbumin inhibitory neurons, RRID:IMSR_JAX:008069), *Sst-Cre* (somatostatin inhibitory neurons, RRID:IMSR_JAX:028864), and *VipCre* (vasoactive intestinal peptide inhibitory neurons, RRID:IMSR_JAX:010908).

### Generation of the Scn1a-flox(R1636Q) mouse

The *Scn1a*-flox(R1636Q) mouse was generated using CRISPR/Cas9 on C57Bl/6J embryos. The allele was designed to insert an additional copy of the last exon of *Scn1a* (exon 27; **Fig 1A**), which includes part of the coding region as well as the 3’ UTR. The upstream copy, which is flanked by loxP sides, encodes arginine at codon 1636 and the bovine growth hormone 3’ UTR. The downstream copy encodes glutamine at codon 1636 (c.4907G>A) and the endogenous 3’ UTR. In the absence of Cre, the upstream copy of exon 27 is spliced into the transcript. After Cre-mediated recombination, the downstream, mutant copy is included. This allele design was based on (20).

**Fig. 1.**
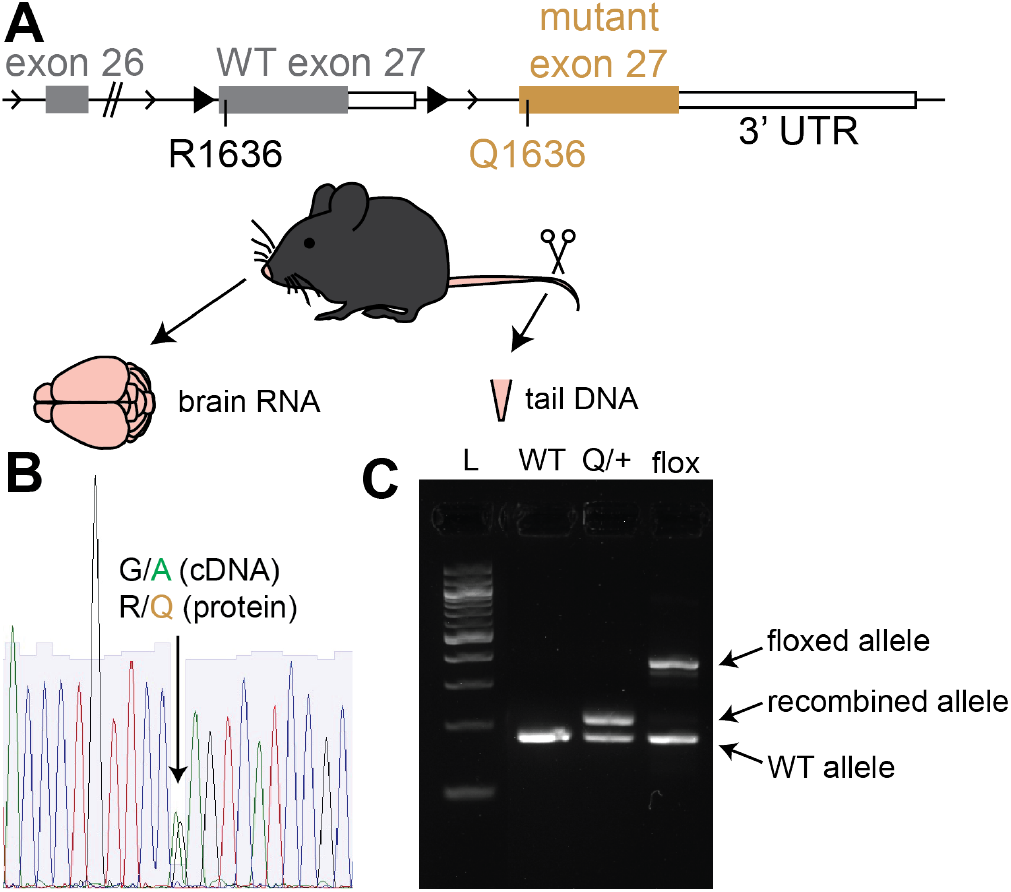
Design and validation of the *Scn1a*-flox(R1636Q) allele. **A**, Structure of the *Scn1a*-flox(R1636Q) allele. The final exon of *Scn1a* was duplicated. The upstream copy, which encodes WT exon 27 followed by the bovine growth hormone 3’UTR, is flanked by loxP sites and is spliced into the mRNA in the absence of Cre. In the presence of Cre, the upstream copy of exon 27 is excised and the mRNA includes the downstream copy of exon 27, which encodes the p.R1636Q variant (c.4907G>A) followed by the endogenous 3’UTR. **B**, Sanger sequencing of whole brain cDNA extracted from *Actb-Cre,Scn1a*^*flox(R1636Q)/+*^ mice shows G/A heterozygosity at *Scn1a*-c.4907. **C**, Genotyping gel from tail DNA. L, ladder. Q/+, *Actb-Cre, Scn1a*^*flox(R1636Q)/+*^ mouse. flox, *Scn1a*^*flox(R1636Q)/+*^ mouse.

Founder mice were bred to C57Bl/6J, and N1+ animals were used for all further experiments. Only animals with the correctly targeted allele, no evidence of random integration, and no off-target editing were used to establish the colony. N1 mice were crossed to C57Bl/6J mice, and the offspring were subsequently crossed to *Actb-Cre* for further validation, or to each other to generate homozygous *Scn1aflox(R1636Q)/flox(R1636Q)* breeders.

To validate the *Scn1a* genotype of the *Scn1a*^*flox(R1636Q)/+*^ and *Actb-Cre,Scn1a*^*R1636Q/+*^ (*Actb-Q/+*) mice, tail DNA and whole brain cDNA were prepared (**Fig 1B-C**). The *Scn1a* locus was amplified by PCR using primers F1 5’ GCAGATACTGGAAAATGATTGG, R1 5’ AGGCGGATGTTGAACAGG, and R2 5’ GCAAGAAACATCCCTGTGG. Tail DNA was visualized with agarose gels. As expected, the WT allele yielded a product of 179 bases, the floxed allele yielded a product of 372 bases, and the recombined allele yielded a product of 213 bases (**Fig 1C**). PCR products were purified and DNA sequence was confirmed by Sanger sequencing. Whole brain cDNA was amplified using the same primers and sequenced via Sanger sequencing. As expected, the sequence was identical to the reference *Scn1a* gene, except at position c.4907, which exhibited a 50/50 mix of G/A, consistent with the expected p.R1636Q variant (**Fig 1B**).

### Video monitoring of Q/+ mice

Beginning on P12, dams and all pups in the litter underwent 24h video monitoring (Ethovision XT, Noldus). Videos were analyzed by a trained observer to identify the time of death and evaluate behavior in the preceding minute. Nonlethal behavioral seizures were occasionally observed (**Video S1**).

### Acute video-EEG monitoring of pre-weanling mice

Seizure-induced mortality in *Actb-Q/+* mice occurs before weaning, and the young mice cannot be separated from the dam for longer than a few hours. Additionally, the small size of the mice prevented implantation of wireless EEG transponders. EEG was performed acutely using a Ponemah wireless EEG system (Data Sciences International).

On P14 or P15, *Actb-Q/+* mice were anesthetized with isoflurane and placed in a stereotaxic instrument. Analgesia (meloxicam 5 mg/kg subcutaneous) was administered. Hair was removed with depilatory cream and the scalp was cleaned with ethanol and betadine. A midline incision was made to expose the skull. Four small holes were drilled: one each over bilateral sensory and motor cortex. Electrodes were placed and secured with dental cement. The incision was closed around the EEG electrodes with sutures. The wireless EEG transponder was suspended from the side of the enclosure.

Mice were allowed to recover on a heating pad until they regained mobility (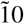 mins), when they were removed to a small, clear-walled enclosure. Video and EEG recording began immediately and continued for up to 8 hours (or until death).

EEG signal was acquired at 500 Hz. EEG data were analyzed using Neuroscore (DSI) software. First, EEG signals were preprocessed by filtering with a powerline filter (60 Hz notch filter) followed by 1 Hz high pass filtering. Then, an analysis protocol for spike and spike train detection was applied to determine periods of abnormal EEG. Spikes were detected when the EEG signal had a minimum amplitude of 200 µV and was greater than the root-mean-squared value of the activity within the preceding minute. Myoclonic jerks were defined as polyspike bursts fast, high-amplitude sharp waves of 0.1 - 0.2s duration correlated with a physical twitch. Seizures were defined as spike trains of at least 5 spikes lasting at least 3s with mean inter-spike interval of 0.05 - 0.6s. Behavior was visually confirmed for all putative events by an expert reviewer.

### Continuous video-EEG monitoring of adult mice

In adult animals, EEG implantation was performed as described (21). Briefly, mice were anesthetized with isoflurane. Four burr holes were drilled through the skull over left and right motor and sensory cortices. Wired leads from a DSI telemetry device (HD-X02) were placed through these holes and secured using dental cement. The corresponding telemetry device was implanted subcutaneously in the back. Animals were allowed to recover for 3 days before EEG recording. Acquisition and data analysis were performed as described above.

### Widefield calcium and optical intrinsic imaging

Experimental mice were generated by crossing *Scn1a*^*flox(R1636Q)/+*^,*Thy1-jRGECO1a* (RRID:IMSR_JAX:030526) mice to *Actb-Cre* mice. On P14 or 15, cranial windows were implanted as described (22) using subcutaneous lidocaine analgesia instead of general anesthesia to avoid contamination of same-day imaging signal. Widefield imaging of calcium signal (jRGECO) was performed on a Leica M205FA stereoscope as described (22). Spontaneous awake brain activity was monitored continuously immediately following cranial window implantation until spontaneous death of animals (typically within 1-4 hours). Data were processed using open-source tools (23), segmenting images using a manually drawn brain mask followed by affine transformation to a common atlas space, background subtraction, and calculation of %dF/F by subtracting and dividing the 20th percentile value from the first 5 seconds of the recording. Temporal bounds of seizure and cortical spreading depolarization (CSD) events were segmented using inflection points in the derivative of the root mean square of the %dF/F signal and verified by visual scoring by a trained clinician.

### Assessment of developmental milestones in Q/+ mice

Developmental milestones were assessed every other day beginning at P4 and ending at P14 (or death). Assays were performed as described (24), and included measurements of body length, tail length, and weight; age at fur appearance, eye and ear opening, incisor eruption, head and forelimb elevation, and quadruped walking; assessment of the surface righting reflex, the tactile reflex, the auditory reflex, the grasp reflex, negative geotaxis, cliff avoidance, and hindlimb clasping, and the ability to climb horizontal and vertical screens. All experiments were performed blind to genotype.

### Audiogenic seizure induction

Adult mice (18-24 weeks old) were placed in a clear plastic arena covered in wire mesh. To induce audiogenic seizures, a sonicator filled with water was turned on for 30 seconds. Mouse behavior was filmed and scored by expert observers for signs of seizure, including wild running/jumping and falling.

### GS967 treatment

GS967 was formulated into mouse chow (Open Standard Diet, Research Diets, 8 mg/kg GS967) (25). *Dlx5/6-Q/+* mice were weaned onto GS967 chow at P21 and maintained on that diet until P60. Survival was monitored daily.

## Results

### Seizures and premature lethality in the Scn1a-p.R1636Q mouse

We generated a mouse with conditional expression of the *Scn1a*-p.R1636Q variant (10) (**Fig 1**), a recurrent variant observed in at least 12 unrelated patients and which causes *SCN1A* GoF epilepsy typically presenting with epileptic encephalopathy and movement disorder (10, 11). Crossing the *Scn1a*^*flox(R1636Q)/+*^ mouse to the globally expressed *Actb-Cre* (**Fig 2A**) yielded the expected Mendelian ratio of *Actb-Cre,Scn1a*^*R1636Q/+*^ (*Actb-Q/+*) pups and WT littermates (*Actb-WT*; 224/487 *Actb-Q/+* vs 263/487 *Actb-WT*; p = 0.0850, Fisher’s exact test; **Fig 2B**), suggesting that the *Scn1a-p.R1636Q* variant does not cause appreciable prenatal lethality in mice. *Actb-Q/+* mice also gained weight at the same rate as their WT littermates (**Fig 2D**). However, 100% (64/64) of *Actb-Q/+* mice died prematurely between P12-18 (**Fig 2C**).

**Fig. 2.**
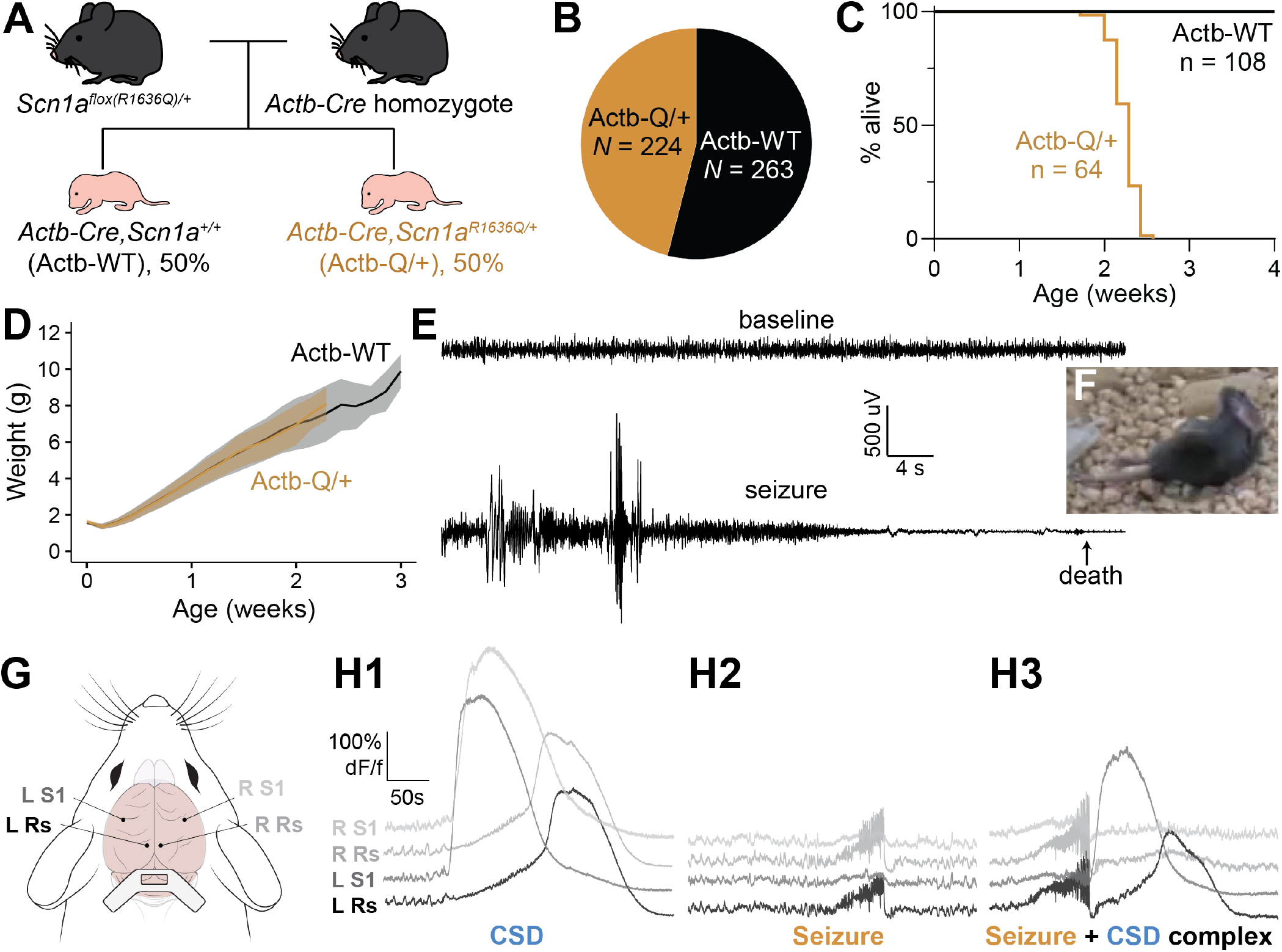
Premature mortality of *Actb-Q/+* mice due to spontaneous seizures. **A**, Breeding scheme to generate *Actb-Q/+* mice. **B**, Proportion of *Actb-WT* vs. *Actb-Q/+* mice at P7 generated by the cross in A (*p* = 0.0850, Fisher’s exact test, measured at P6-7). **C**, Survival of *Actb-WT* and *Actb-Q/+* mice (*p* < 0.0001, Mantel-Cox log-rank test). **D**, Weight of *Actb-WT* and *Actb-Q/+* mice (genotype *p* = 0.3410, two-way ANOVA). **E**, EEG of a P14 *Actb-Q/+* mouse. Top, baseline EEG. Bottom, EEG during a lethal seizure. **F**, Seizure posture includes hindlimb extension. **G**, Configuration for widefield imaging in P14-15 pups (S1, primary somatosensory cortex; Rs, retrosplenial cortex). **H**, Example widefield calcium imaging traces showing the three abnormal phenomena observed in P15 *Actb-Q/+* mice: isolated cortical spreading depolarizations (CSDs), isolated seizures, and CSD + seizure complexes.

To investigate the cause of death, we performed 24-hour video monitoring of the dam and litter starting at P12. We observed behavioral seizures immediately preceding death in every *Actb-Q/+* mouse (13/13 mice), and several instances of apparent behavioral seizures that did not cause death (**Video S1**). To confirm these events were seizures, we performed video-EEG monitoring in *Actb-Q/+* mice on P14 or P15. EEG implantation and recordings were performed acutely, and recordings lasted 8-10 hours, which facilitated capture of a lethal event in an *Actb-Q/+* mouse (**Fig 2E**), which consisted of behavioral seizure (**Fig 2F**) accompanied by large-amplitude spikes on EEG and culminating in hind limb extension and death.

The EEG implantation procedure requires prior anesthesia, which may have artificially suppressed seizure activity in subsequent recording sessions. Therefore, we further confirmed spontaneous seizure activity using *in vivo* widefield calcium imaging, which allowed us to capture brain activity simultaneously across the entire dorsal brain surface (**Fig 2G-H**). In *Actb-Q/+* mice, but not WT littermates, we observed several abnormal phenomena. *Actb-Q/+* mice exhibit putative spontaneous seizures (**Fig 2H2**) behaviorally similar to events captured on video monitoring and video EEG and that consisted of diffuse rhythmic bursting lasting 40-150 seconds. These seizures often trigger post-ictal cortical spreading depolarization (CSDs; **Fig 2H3**), slow-moving waves of ionic disruption which are linked to migraine aura and which may participate in seizure termination (26) and which have also been observed in *Scn1a*^*+/-*^ mice (27). We also observed isolated CSDs occurring in the apparent absence of seizures (**Fig 2H1**).

### Global expression of Scn1a-p.R1636Q causes motor dysfunction in developing mice

The human phenotype associated with *SCN1A*-p.R1636Q includes epilepsy and global developmental delay, as well as a prominent movement disorder which emerges around 2 years of age and includes chorea, dystonia, myoclonus, and/or ataxia (10, 11). To assess whether *Actb-Q/+* mice exhibit developmental delay or motor dysfunction, we performed a battery of developmental milestones, including several assessments of motor function, in *Actb-Q/+* mice and their WT littermates every other day between P4 – P14 (21, 24).

All assessments of physical development (body length, fur appearance, eye opening, ear opening, etc.) were normal in *Actb-Q/+* mice (**Fig 3**). However, *Actb-Q/+* mice exhibit reduced completion of the vertical screen test at P10 (*Actb-WT*: 3/51 mice successfully competed; *Actb-Q/+*: 0/43 mice successfully completed; *p* = 0.0637, Fisher’s exact test), P12 (*Actb-WT*: 18/64 mice successfully completed; *Actb-Q/+*: 4/55 mice successfully completed; *p* = 0.0017, Fisher’s exact test), and P14 (*Actb-WT*: 45/56 mice successfully completed; *Actb-Q/+*: 26/43 mice successfully completed; *p* = 0.0423, Fisher’s exact test; Fig 3), suggesting that *Actb-Q/+* mice have mild impairments in strength and/or coordination (28).

**Fig. 3.**
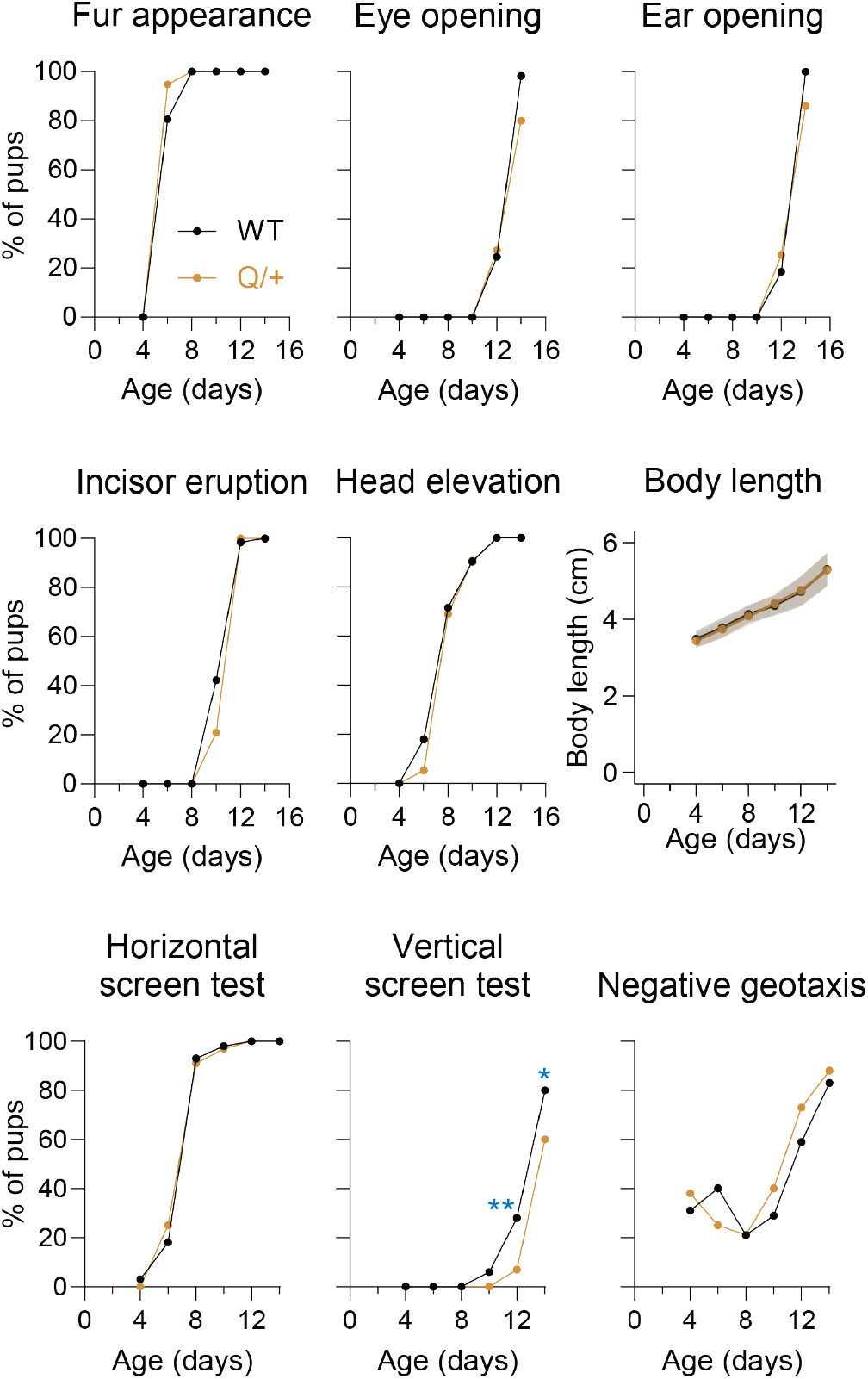
*Actb-Q/+* mice exhibit mild motor impairment during development. **A**, *Actb-Q/* + mice and WT littermates were subjected to a battery of developmental behavioral tests every other day between P4 and P14. * indicates *p* < 0.05; ** indicates *p* < 0.01. Statistical analysis performed via Fisher’s exact tests. For each age and genotype, *N* = 37-61 mice.

### Dysfunction of developing parvalbumin interneurons causes seizures and premature mortality in Scn1a-p.R1636Q mice

To determine which cell type(s) might be responsible for the *SCN1A* GoF epilepsy phenotype, we crossed the *Scn1a*^*flox(R1636Q)/+*^ mouse to a series of cell type-specific Cre driver lines. Activation of the *Scn1a*-p.R1636Q allele in excitatory neurons with *Slc17a7-Cre* did not cause premature lethality or any observable deficit (**Fig 4A**). We next tested the sufficiency of activating the *Scn1a*-p.R1636Q allele in all inhibitory neurons using *Dlx5/6-Cre*, which is expressed in all inhibitory neuron progenitors in both the medial and caudal ganglionic eminences (29, 30). We found that 100% of *Dlx5/6-Q/+* mice die between P25 and P33 (10/10 mice; **Fig 4A**). To identify the cause of death in these mice, we performed 24-hour video monitoring in 5 *Dlx5/6-Q/+* mice. All 5 mice died immediately following a convulsive seizure. Non-lethal seizures were also observed. These data suggest that inhibitory neuron dysfunction is the primary driver of disease in *SCN1A* GoF epilepsy.

**Fig. 4.**
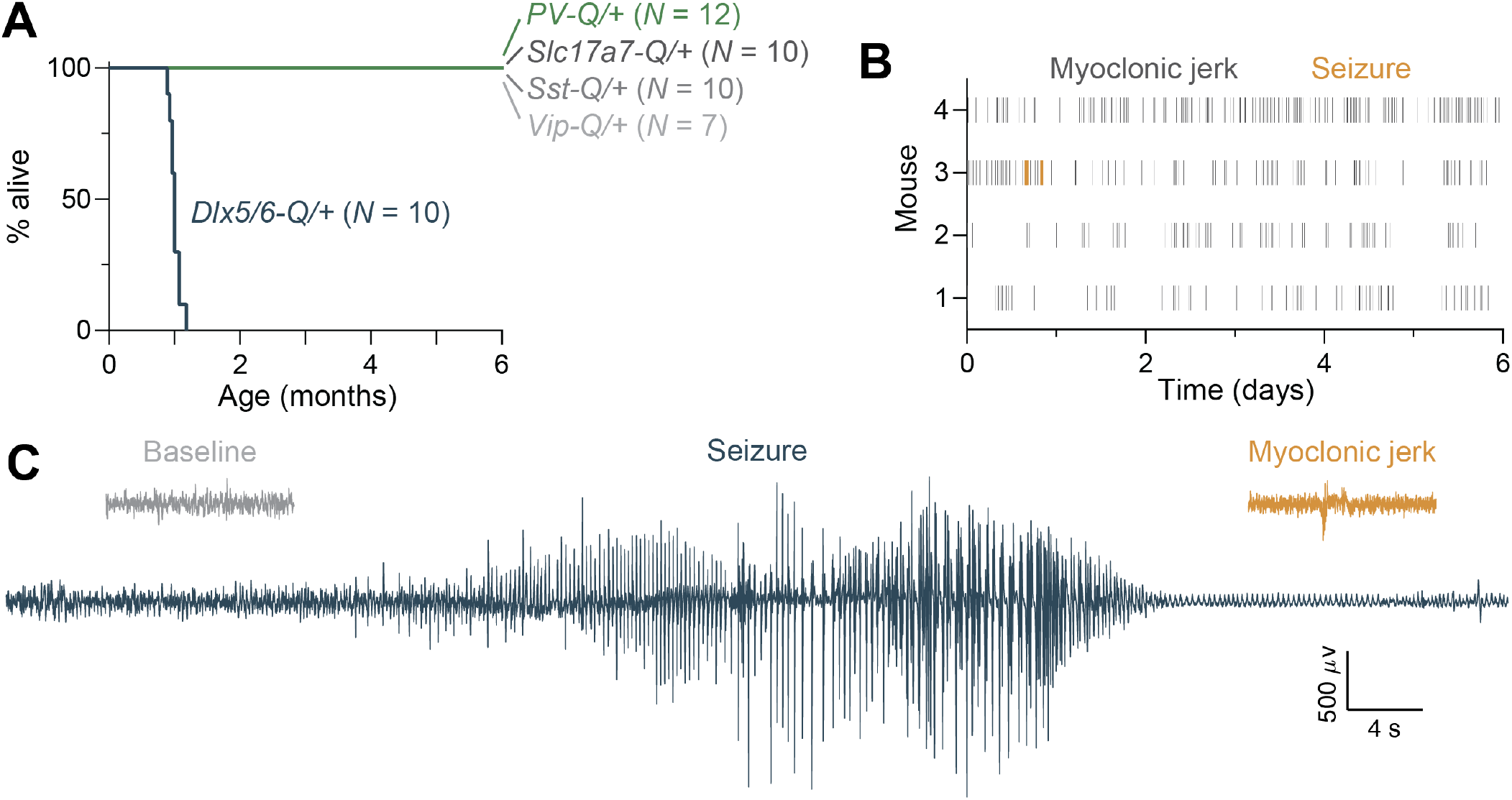
Activation of the *Scn1a*-R1636 allele in inhibitory neurons largely recapitulates the global phenotype. **A**, Survival of cell type-specific *Scn1a*^*flox(R1636Q)/+*^ mice. **B**, Myoclonic jerk and seizure frequency in 4 *PV-Q/+* mice monitored via EEG for 6 days. **C**, Example baseline, myoclonic jerk, and seizure EEG traces from a *PV-Q/+* animal.

We attempted to narrow down the inhibitory neuron population(s) involved by activating the *Scn1a*-p.R1636Q allele specifically in parvalbumin-, somatostatin-, or vasoactive intestinal peptide (VIP)-expressing interneurons. However, crosses to *PV-Cre, Sst-Cre*, and *Vip-Cre* did not reproduce the premature lethality phenotype observed in *Actb-Q/+* and *Dlx5/6-Q/+* mice. No deaths were observed in mice from these crosses at up to 6 months of age (**Fig 4A**).

Despite the lack of mortality, normal handling of *PV-Q/+* mice revealed sound- and vibration-induced behavioral abnormalities, which we hypothesized were related to seizures. To determine whether *PV-Q/+* mice exhibit spontaneous seizures, we performed 6 days of continuous video-EEG monitoring in 4 adult (4 month old) animals. In that time, we observed frequent myoclonic jerks with EEG correlate in 4/4 mice (70.5 ± 80.6 per day; **Fig 4B-C**). One mouse also exhibited 4 spontaneous seizures (**Fig 4B-C**). Taken together, our results suggest that interneurons, and particularly parvalbumin-expressing interneurons, are the primary cell type involved in the pathogenesis of *SCN1A* GoF epilepsy, similar to previous findings in *Scn1a* LoF models (31, 32). However, expression of the *Scn1a*-p.R1636Q variant in developing, Dlx5/6+ interneurons (before parvalbumin itself is expressed, at around P10-14) appears to be required for premature lethality.

### GS967 prolongs lifespan in Scn1a-p.R1636Q mice

We attempted to treat our *SCN1A* GoF epilepsy mice with GS967, a sodium channel blocker that reduces persistent sodium current, enhances entry into the inactivated state, and impairs recovery from inactivation (25, 33–35). We treated *Dlx5/6-Q/+* mice with a GS967 diet from P21 to P60 (**Fig 5**). During the treatment window, only 2/6 *Dlx5/6-Q/+* mouse died, compared to 10/10 untreated mice (*p* = 0.0025, Mantel-Cox log-rank test). After P60, when the GS967 diet was removed, *Dlx5/6-Q/+* mice died rapidly (2/2 mice died within 4 days), supporting the conclusion that treatment with GS967 reduces lethality in this mouse model.

**Fig. 5.**
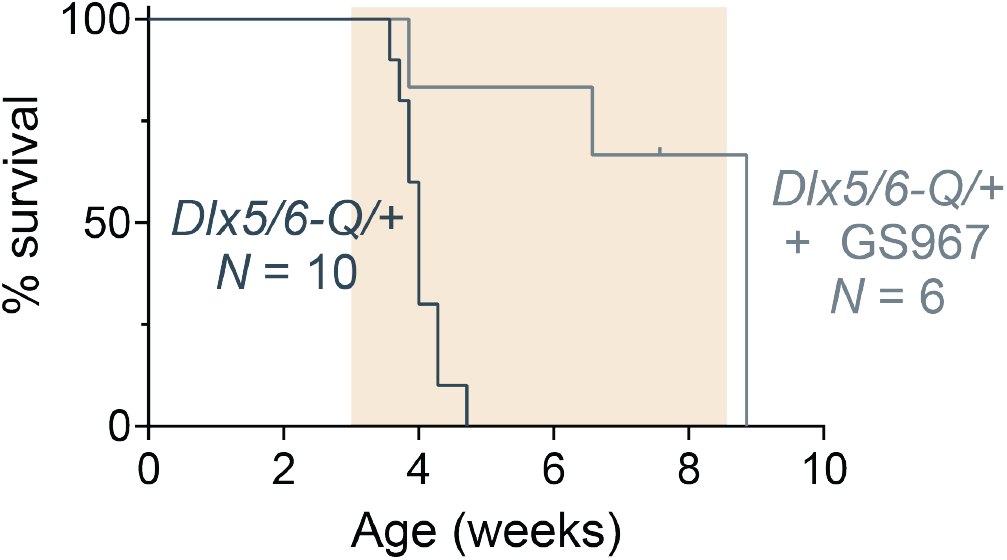
Treatment with GS967 reduces lethality in *Dlx5/6-Q/+* mice. Survival of *Dlx5/6-Q/+* mice fed GS967-supplemented chow from P21 to P60 (indicated by the orange box). Note that mice are rapidly deceased upon removal of GS967.

## Discussion

We developed and characterized the first mouse model of *SCN1A* GoF epilepsy, a recently described disease characterized by early seizure onset (<3 months of age), movement disorder, intellectual disability/developmental delay, and premature mortality (10, 11). Global activation of the *Scn1a*-p.R1636Q variant in mice recapitulates many aspects of the disease, including spontaneous seizures, motor impairment, and premature mortality. Activation of the variant in specific cell types suggests that dysfunction of developing parvalbumin interneurons is primarily responsible for this phenotype. Finally, we demonstrated that modulating sodium channels with GS967 markedly reduces lethality in the *SCN1A* GoF epilepsy mice. Our work is the first mechanistic study of *SCN1A* GoF epilepsy in a preclinical experimental model in vivo. Further investigation in the *Scn1a*-flox(R1636Q) mouse could yield new therapeutic advances and mechanistic insights for the treatment of *SCN1A* GoF epilepsy.

*SCN1A* GoF epilepsy exhibits earlier onset than Dravet syndrome (10, 11). This phenotypic difference was recapitulated in our *Scn1a*-p.R1636Q mouse. Premature lethality in the *Actb-Q/+* mouse begins at P12, and all mice are deceased by P18. In contrast, only 20-50% of *Scn1a*^*+/-*^ mice exhibit premature mortality, and most deaths occur between P18-24 (4, 12, 13). Mortality in *Scn1a*^*+/-*^ mice is heavily dependent on genetic background (12, 36, 37), and future studies of the *Scn1a*^*flox(R1636Q)*^ mouse could assess the effect of different genetic backgrounds on the *SCN1A* GoF epilepsy phenotype. Due to the early mortality in *Actb-Q/+* mice, we were unable to use conventional behavioral testing – typically performed in young adult mice – to evaluate whether *Actb-Q/+* mice exhibit other aspects of the human phenotype such as intellectual disability and features of autism spectrum disorder, which have previously been reported in *Scn1a*^*+/-*^ mice (38). However, we were able to detect mild motor impairment in *Actb-Q/+* mice using a battery of developmental tests used to evaluate juvenile mice.

Yet, it remains unclear how GoF variants can cause both *SCN1A* GoF epilepsy and FHM3. The two diseases appear to be caused by non-overlapping groups of variants, suggesting a genotype-phenotype correlation with an unknown mechanistic basis. Paradoxically, the GoF variants associated with FHM3 tend to cause seemingly more severe effects on channel biophysics than those associated with *SCN1A* GoF epilepsy (10, 11, 17, 18, 39).

While this study reports the first mouse model of *SCN1A* GoF epilepsy, two mouse models of FHM3 have been published to date. *Scn1a*-p.L263V and *Scn1a*-p.L1649Q mice exhibit elevated susceptibility to cortical spreading depolarizations (CSDs) (40–42), which are thought to underlie migraine aura (43). We observed spontaneous CSDs in *Actb-Q/+* mice, both in isolation and as post-ictal events. Post-ictal CSDs may be a natural mechanism for seizure termination (44). In contrast, spontaneous CSDs have also been reported in *Scn1a*^*+/-*^ mice, where they have been hypothesized to contribute to disease pathogenesis (27). Why spontaneous CSDs are observed in mice harboring both GoF and LoF variants in *Scn1a* is also unclear.

We detected seizures across the neocortical surface in *Actb-Q/+* mice, but seizure activity was especially prominent in retrosplenial cortex, a region with strong connections to subcortical structures (especially hippocampus and thalamus) (45). This finding may suggest that seizures in the *ActbQ/+* mice originate sub-cortically, perhaps in subiculum or CA1 hippocampus. Consistent with this interpretation, we occasionally observed near-simultaneous onset of apparently bilateral CSD. Alternatively, the prominent signal in retrosplenial cortex may be an artifact of epifluorescence imaging. The dorsal hippocampus is located directly below retrosplenial cortex, and apparent retrosplenial signal could instead reflect ictal activity in underlying hippocampus. In either scenario, the hippocampus appears to be an important locus of pathology in *SCN1A* GoF epilepsy, at least in mouse.

Dravet syndrome, FHM3, and *SCN1A* GoF epilepsy all appear to be primarily caused by inhibitory neuron dysfunction. The primary cell types affected by *Scn1a* loss are inhibitory neurons expressing parvalbumin (13, 15), somatostatin (31, 38), and vasoactive intestinal peptide (46, 47). Heterozygous deletion of *Scn1a* in parvalbumin interneurons causes spontaneous and temperature-induced seizures as well as premature death (32, 48). Conversely, gene replacement therapies targeted to inhibitory neurons improve *Scn1a*^*+/-*^ mouse phenotypes (49, 50). In the FHM3 model *Scn1a*-p.L1649Q, inhibitory neurons as a group are mildly hyperexcitable (42). Here, we report that expression of the *SCN1A* GoF epilepsy variant *Scn1a*-p.R1636Q in developing inhibitory neurons or specifically in parvalbumin interneurons causes epilepsy, while activation in excitatory neurons is without apparent phenotype. Yet, the underlying mechanistic basis of the stark differences between distinct *SCN1A* spectrum disorders remains unclear. It is possible that variants associated with FHM3, typically more severe at the level of the ion channel, induce hyperexcitability of PV+INs, while variants associated with *SCN1A* GoF epilepsy with apparently more mild GoF effects at the channel level instead lead to impaired excitability of PV+INs akin to or perhaps even more severe than observed with LoF/in Dravet syndrome.

Our findings shed light on the necessity of Na_V_1.1 for interneuron development and the potential role of Na_V_1.1 in pyramidal cells in the pathogenesis of *SCN1A* spectrum disorders. Expression of the variant in interneuron precursor cells beginning around E12.5 (29) with *Dlx5/6-Cre* resulted in a severe phenotype mimicking the global mutant. However, expression of the variant in post-mitotic, post-migration parvalbumin interneurons at/around P14 and likely beginning no earlier than P10 using *PV-Cre* results in a comparatively milder phenotype, including myoclonic jerks and rare spontaneous seizures but without premature lethality. Expression in excitatory cells yielded no apparent phenotype.

In the clinic, sodium channel blockers can be partially effective for the treatment of epilepsy in patients with *SCN1A* GoF syndromes (10, 11). Our prior work showed partial normalization of currents mediated by variant Na_V_1.1-p.R1636Q subunits and seizure suppression in a human patient with oxcarbazepine (10). Here we show this response to sodium channel blockers was also recapitulated in our mouse model. *Dlx5/6-Q/+* mice were protected from premature death while treated with GS967, a sodium current blocker. This finding supports the future utility of the *Scn1a*^*flox(R1636Q)*^ mouse for screening of potential therapeutics to treat *SCN1A* GoF epilepsy.

## ACKNOWLEDGEMENTS

This work was supported by R01 NS110869 (EMG), the Dravet Syndrome Foundation Research Grant (EMG), the Dravet Syndrome Foundation Postdoctoral Fellowship (SFH), the Brody Family Medical Trust Fellowship in Incurable Diseases (SFH), the Holt Family Epilepsy Neurogenetics Fellowship (SFH), and the Jump-Start Research Tools Matching Grant Program Award from the University of Pennsylvania Orphan Disease Center (EMG). We gratefully acknowledge technical support provided by Dr. Xiaohong Zhang and Clara Wang. We also thank the *SCN1A* Gain of Function Foundation for participation in our clinical research.

